# Contrasting hierarchical and multiple-demand accounts of frontal lobe functional organisation during task-switching

**DOI:** 10.1101/2020.06.27.175133

**Authors:** Richard E. Daws, Eyal Soreq, Yuqi Li, Stefano Sandrone, Adam Hampshire

## Abstract

There is an unresolved discrepancy between popular hierarchical and multiple-demand perspectives on the functional organisation of the human frontal lobes. Here, we tested alternative predictions of these perspectives with a novel fMRI switching paradigm. Each trial involved switching attention between stimuli, but at different levels of difficulty and abstraction. As expected, increasing response times were evident when comparing low-level perceptual switching to more abstract dimension, rule and task-switching. However, there was no evidence of an abstraction hierarchy within the prefrontal cortex (PFC). Nor was there recruitment of additional anterior PFC regions under increased switching demand. Instead, switching activated a widespread network of frontoparietal and cerebellar regions. Critically, the activity within PFC sub-regions uniformly increased with behavioural switch costs. We propose that both perspectives have some validity, but neither is complete. Too many studies have reported dissociations within MD for this volume to be functionally uniform, and the recruitment of more anterior regions with increased general difficulty cannot explain those results. Conversely, whilst reproducible evidence for a hierarchical functional organisation has been reported, this cannot be explained in terms of abstraction of representation or reconfiguration *per se*, because those interpretations generalise poorly to other task contexts.

## Introduction

The prefrontal cortex (PFC) is critical for human cognition and behaviour (Shallice and Evans, 1978; Fuster, 1999; Duncan, 2001), yet how regions within the PFC are functionally organised remains controversial. Broadly speaking, much of the debate is polarised around two competing perspectives.

One prominent class of models revolve around the notion that the PFC is organised hierarchically (Figure 1), with increasingly anterior PFC (aPFC) regions placing towards the top of the hierarchy and supporting cognitive processes of an increasingly higher-order (Koechlin, Ody and Kouneiher, 2003; Ramnani and Owen, 2004; Badre, 2008; Badre and D’Esposito, 2009). Evidence in support of this hypothesis has been provided from studies of executive dysfunction after lesions to aPFC (Nelson, 1976; Reitan and Wolfson, 1994; Duncan, Burgess and Emslie, 1995; Shallice *et al.*, 2007; Badre *et al.*, 2009; Roca *et al.*, 2010), and by findings of increased functional MRI (fMRI) aPFC activation for tasks that require switching (Koechlin *et al.*, 1999; Rushworth, Passingham and Nobre, 2002; Braver, Reynolds and Donaldson, 2003; Koechlin, Ody and Kouneiher, 2003), sequential processes (Koechlin *et al.*, 1999; Vendetti and Bunge, 2014), higher-order integrations (Parkin *et al.*, 2015) and reversal learning (Hampshire and Owen, 2006).

**Figure 1.**
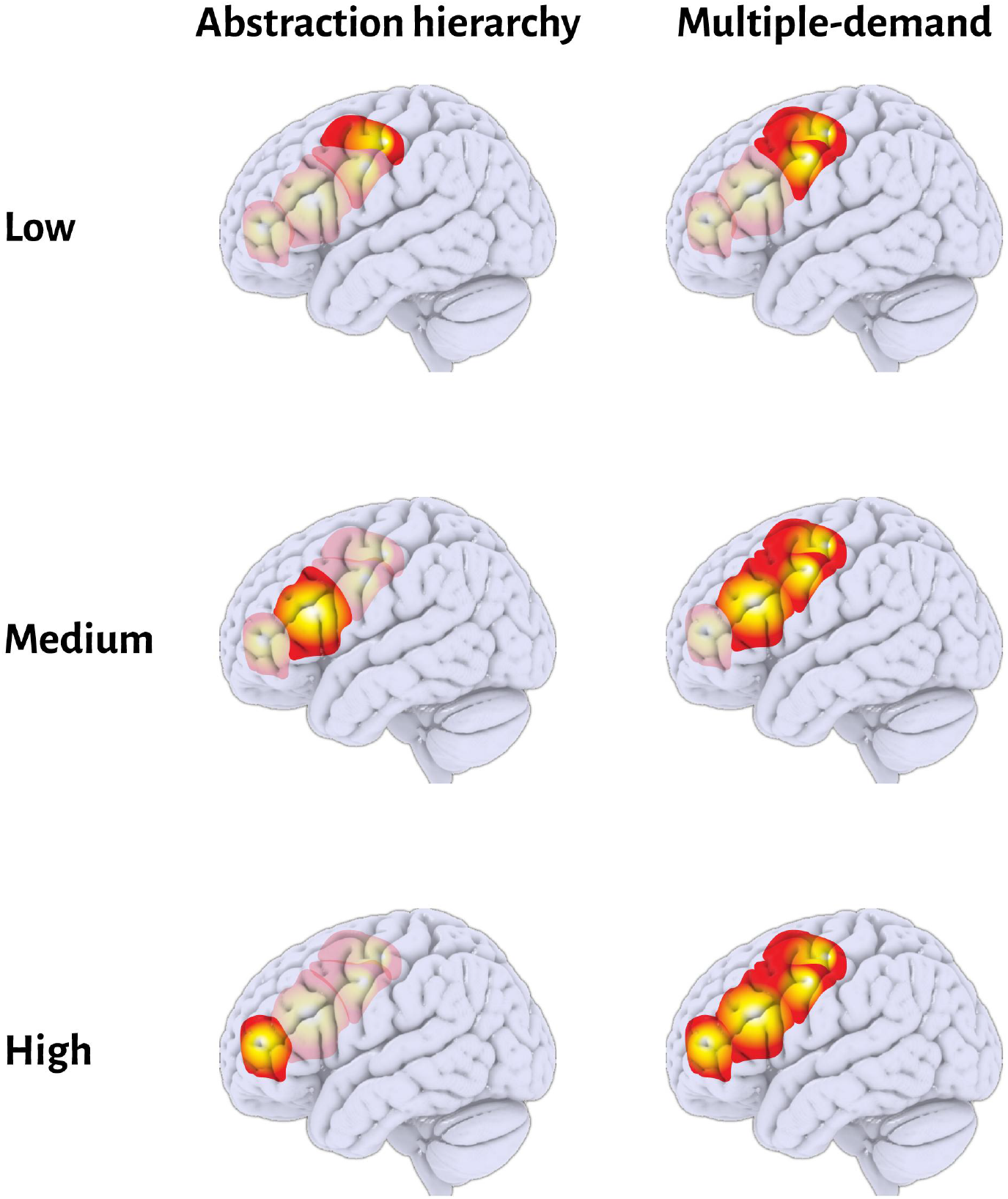
Schematic of alternate accounts of the frontal lobe’s functional organisation. Left column depicts hierarchical models (Koechlin et al. 2003; Badre 2008) that denote increasingly anterior prefrontal cortex (aPFC) regions placing towards the top of the hierarchy and supporting cognitive processes of an increasingly higher-order (top-to-bottom rows). The right column depicts an alternate multiple-demand cortex (MDC) model (Duncan 2013; Crittenden and Duncan 2014). Here, MDC forms a distributed cognitive resource that flexibly recruits additional, and more anterior, frontal regions under increased task difficulty (top-to-bottom rows).

However, there is increasing awareness that the generalisation of one-to-one mappings between cognitive processes and regional activations is highly problematic. More specifically, the same brain regions are often activated during tasks that operationally are quite distinct (Duncan and Owen, 2000; Duncan, 2013; Fedorenko, Duncan and Kanwisher, 2013), whilst brain regions attributed specific functions are rarely activated in isolation (Daws et al. 2020; Hampshire et al. 2010; Mouraux et al. 2011; Yarkoni et al. 2011; Hampshire and Sharp 2015; Lorenz et al. 2017).

An alternative perspective that can account for this general activation profile states that a set of brain regions, referred to as “multiple-demand” cortex (MDC), forms a flexible cognitive resource (Duncan and Owen, 2000; Fedorenko, Duncan and Kanwisher, 2013). MDC is proposed to be recruited during diverse demanding tasks. In support of this view, aPFC, is often activated as part of a broader distributed network during diverse task contexts. Task-switching is one example (Dove et al. 2000; Cusack et al. 2010; Daws et al. 2020), but others include target detection (Hampshire, Duncan and Owen, 2007), motor-inhibition (Erika-Florence, Leech and Hampshire, 2014) and instruction-based learning (Erika-Florence, Leech and Hampshire, 2014; Hampshire *et al.*, 2016, 2019). aPFC activation has also been reported during tasks that lack abstraction or hierarchy by design, but that are challenging in other ways (Jiang and Kanwisher, 2003; Crittenden and Duncan, 2014). In order to account for studies that appeared to support PFC hierarchy, proponents of the globalist perspective (Dehaene et al. 1998) have proposed that progressively larger, and more anterior portions of the frontal lobes, including aPFC, may be recruited to the MDC network under increased task difficulty in general (Jiang and Kanwisher, 2003; Crittenden and Duncan, 2014).

In this study, we developed a ‘multilevel switching’ (ML) paradigm to test the alternative predictions of the hierarchical and MDC perspectives on PFC organisation. The aims of the ML paradigm were to compare switching conditions that differentiate the level of abstraction in information that must be updated during the switch from general task difficulty, as gauged by the relative switch cost on response time.

According to the hierarchical perspectives, there should be a tight mapping of switch abstraction to PFC regions, ranging from low-level perceptual changes, through visual to dimension, mapping rule and task-switching, and with task-switching selectively activating aPFC. Conversely, the multiple-demands account would predict more PFC brain regions being recruited along an anterior-posterior gradient as the magnitude of the behavioural switching cost increases, that is, regardless of the level of abstraction.

## Methods

### Participants

15 young healthy adults (8 female, mean age 25, ranging 20-40 years) participated in the study. All participants were right-handed English speakers with normal, or corrected to normal, eyesight. Volunteers were excluded if they had a history of neurological or psychiatric illness, were taking psychoactive medications or did not meet MRI safety criteria. Approval for this study was received by the Cambridge university research ethics committee and participants gave informed consent prior to entering the fMRI scanner.

### Multilevel switching task design

We designed the ML switching paradigm (Figure 2a) to compare attentional switches that varied in difficulty, gauged by behavioural switch costs, and level of abstraction of information that must be updated during the switch. Each trial presented a target and individual’s identified which of two concurrently presented probes was correct based on the current mapping rule. At the highest level, switches could occur between two *tasks* that involved processing arrays of either words or coloured shapes. Within each task, switches could occur between two available discrimination *rules.* Rules related to whether probes were being selected based on which probe *matched* the target, or which probe was the *odd one out*, relative to the target and the other probe. Switches also could occur between one of two *dimensions* of the stimuli within each task. For words, these dimensions were the physical size of the described object or animate/inanimate. For objects, the dimensions were shape and colour. At the lower perceptual levels, there were *comparison* switches, where probe exemplars switched for one of the dimensions, *target* switching, where the target stimulus was updated but probes remained the same, and *side* switches, where the two probes switched positions and target remained the same, requiring a different motor response. One of these 6 switches, task, rule, dimension, target, comparison or side, occurred on every trial. Notably, rule dimension, and target switches were designed to be similar insofar as they all differ by just one aspect to the comparison switch, and two aspects to each other.

**Figure 2.**
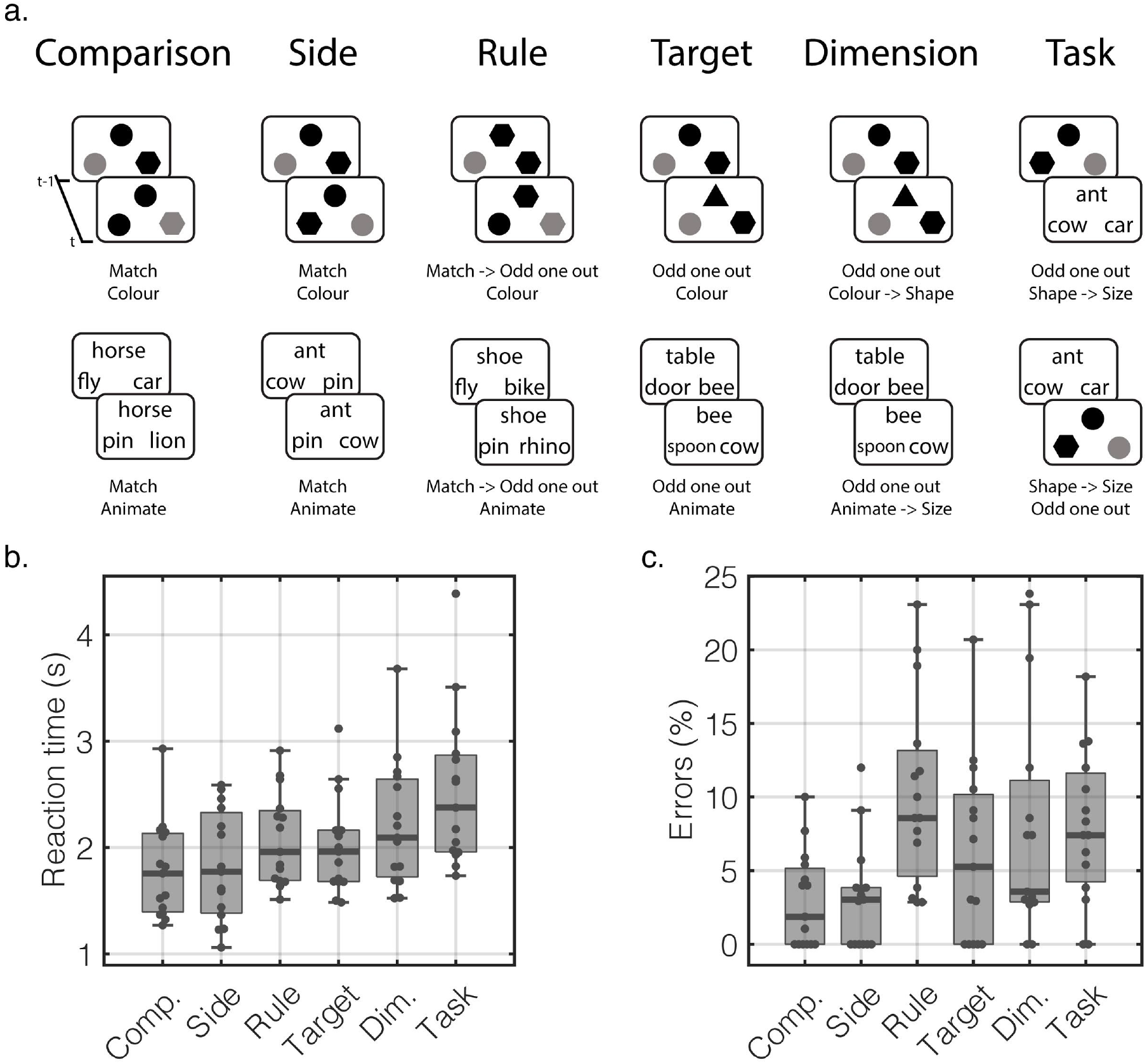
Multilevel switching (ML) paradigm switch costs. a) Each trial (t) involved a switch relative to the previous trial (t-1). Left or right responses were made, relative to the target and depending on the current rule and dimension. Examples of switches from the object and word tasks on the top and bottom rows, respectively. b) Individuals median reaction time in seconds (s) for correct responses in each condition (ranked by the group-level median). c) The percentage of response errors made for each condition.

### Data acquisition & preprocessing

Responses were made on an MRI compatible button box using the 1st and 2nd fingers of the right-hand. Tasks were programmed in Visual Basic and stimuli were projected on a screen, visible via a mirror, at the end of the scanner bore. Brain images were collected using a 3 Tesla Siemens Scanner. A T2-weighted echo planar image depicting blood oxygenation level dependent (BOLD) contrast was acquired every 2s. The first 10 images were discarded to account for equilibrium effects. Images consisted of 32 * 3 mm slices, with a 64 × 64 matrix, 192 × 192 mm field of view, 30 ms TE, 2 s TR, 78° flip angle, 0.51 ms echo spacing, and 2232 Hz/Px bandwidth. A 1 mm resolution MPRAGE T1-weighted structural scan was also collected for each individual with a 256 × 240 × 192 matrix, 900 ms TI, 2.99 ms TE and 9° flip angle. Data were pre-processed using a standard pipeline in SPM12 (Statistical Parametric Mapping, Welcome Department of Imaging Neuroscience). Specifically, images were slice-timing and motion-corrected, spatially warped onto the standard Montreal Neurological Institute template using the structural scan, and spatially smoothed with an 8 mm full width at half maximum Gaussian kernel.

### Univariate analysis

Individual’s fMRI data were modelled using voxelwise general linear models (GLMs) in SPM12. Psychological predictors for each of the 6 switch types, defined from trial onsets and durations until response, were convolved with the canonical haemodynamic response function. Head motion was modelled using 12 parameters derived from the rigid-body motion-correction stage concatenated with their 1st-order temporal derivatives. Individual’s beta maps were taken to the group-level and modelled within a repeated-measures analysis of variance (rm-anova) to examine switching (6 levels). Unless otherwise stated, group-level activation maps are initially thresholded at p<0.01 uncorrected, followed by a cluster-level false discovery rate (FDR) p<0.05 correction for multiple-comparisons across the whole brain.

### Region of interest definition

Our in-house developed watershed transform (Grant *et al.*, 2018; Soreq, Leech and Hampshire, 2019; Hampshire *et al.*, 2020) was used to segment functional activation maps into discrete clusters in a data-driven manner. This common segmentation considers a 3D statistical volume (e.g., activation map) as a multi-dimensional surface where high and low intensities represent elevations. The algorithm iteratively “fills” independent catchment basins (CB) with unique labels by flooding the various-independent local minima in the statistical volume and their surroundings. This results in the continuous statistical space of the activation map being converted into a discrete set of regions of interest (ROI) that can be non-gaussian in shape, whilst accounting for contiguous functional regions that have multiple maxima.

## Results

### Behavioural performance relates to switch abstraction

As expected based on prior behavioural pilotting, median reaction times (RT) for correct responses varied across the switching conditions (one-way rm-anova: F(5, 70)=20.265, p<0.001, 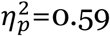) and the pairwise comparisons (Figure 2b) partially differentiated switching abstraction from global difficulty as gauged by the behavioural switch cost. Specifically, the group-level RT was slowest for task-switches (median=2.375, interquartile range (IQR)=0.910), and was significantly slower than dimension switches (paired t-test: t_14_=3.280, p=0.014, confidence interval 95% (CI)=0.11 to 0.53, *d*=0.85), which was significantly slower than target switches (t_14_=2.525, p=0.041, CI=0.03 to 0.36, *d*=0.65), in turn. Target and rule switch RT did not differ (t_14_=1.031, p=0.320, CI=-0.18 to 0.06, *d*=0.27). RT for Rule switching was significantly slower than side switching (t_14_=4.330, p=0.004, CI=0.13 to 0.38, *d*=1.12), and side and comparison switch RT did not differ (t_14_=0.625, p=0.542, CI=−0.10 to 0.18, Cohen’s *d*=0.16) (all p-values FDR-corrected).

Therefore, using RT as a behavioural index of cognitive demand controls general difficulty whilst probing switch abstraction across the target and rule conditions. It also provides a broader difficulty axis based on response speed, with low-level perceptual comparisons and side switches being the most simple, target, dimension and rule switches having medium difficulty, and task switches being the most demanding.

Response accuracy (Figure 2c) for all trials was near ceiling (median=95.802%, IQR=7.222), but significantly varied across switch type (one-way rm-anova: F(5,70)=6.712, p<0.001, 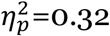). Response errors were most frequent for the rule (median=8.571%, IQR=8.560) and task (median=7.407%, IQR=7.400) switches, and these types did not differ in frequency (paired t-test: t_14_=1.606, CI=−0.01 to 0.05, p=0.131, *d*=0.42). Rule switching errors were significantly more common than target switch errors (t_14_=3.288, CI=0.01 to 0.07, p=0.005, *d*=0.85), and there was no difference between task and target switch errors (t_14_=1.153, p=0.268, CI=−0.02 to 0.05)

### Switching activates a broad network of brain regions

Activation during switching was modelled in a one-way rm-ANOVA (6-levels). Collapsing across switch types, switching increased activation across a broad set of lateral frontoparietal, anterior insula, occipital and cerebellar regions (Figure 3a). The inverse contrast rendered activation decreases in ventromedial PFC (vmPFC), posterior cingulate cortex (PCC) and in the temporal parietal junction (TPJ). This accords with an increase in MDC and reduction in DMN activity during stimulus-driven switching (Daws et al. 2020).

**Figure 3.**
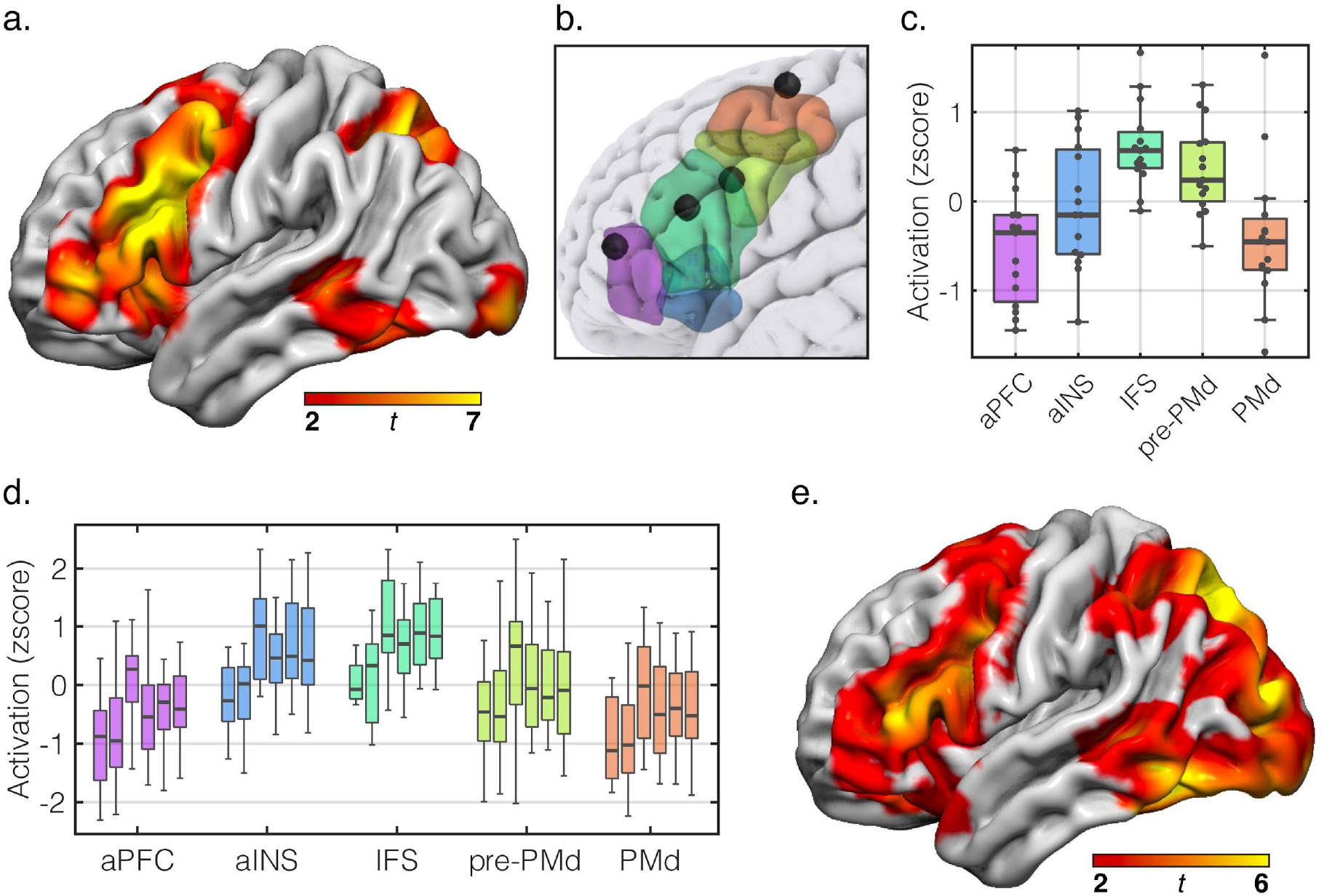
Uniform activation increases in lateral frontal cortex (LFC) with increasing switching demands. a) Activation increases associated with switching, in general. b) LFC ROIs derived from a watershed transform of the switching activation map (a). Coordinates from the rule abstraction hierarchy (Badre and D’Esposito, 2007) are rendered as 5mm radius black sphere overlays. c) ROI mean activation across switching conditions (z-scored within subject). d) ROI mean activation for each ROI and condition (left-to-right: comparison, side, rule, target, dimension, task). e) Supplemental voxelwise t-contrast testing the effect of cognitive demand, indexed by RT, on activation. All maps are thresholded at p<0.01 uncorrected, followed by a p<0.05 cluster-correction. (aPFC=anterior prefrontal cortex, aINS=anterior insula, IFS=inferior frontal sulcus, PMd=dorsal premotor cortex).

### PFC activation during switching does not reveal a hierarchy

Focused ROI analyses were applied to further test for evidence of a posterior-posterior hierarchy. We segmented the activation map for the contrast of all switches (i.e., average contrast value relative to implicit baseline) using a watershed transform (see methods). This separated switching activation in the left frontal lobe into 5 ROIs (Figure 3b) that closely aligned with the proposed coordinates of the rule abstraction hierarchy (Badre and D’Esposito, 2007) and the information cascade model (Koechlin et al. 2003): dorsal premotor (PMd), pre-PMd, inferior frontal sulcus (IFS), aPFC and the anterior insula (aINS). We used the group-level statistical map to extract a weighted-mean beta activation from each ROI that was then z-scored within-individual and compared in a two-way rm-anova with ROI (5 levels) and switch type (6 levels) as within subject factors. There were significant main effects of ROI (F(4, 56)=7.175 p<0.001, 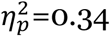) and switch type (F(5, 70)=7.190, p<0.001, 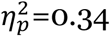). Critically though, there was no ROI * switch type interaction (F(20, 280)=0.960, p=0.450, 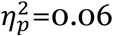) (p-values Greenhouse-Geisser adjusted).

The main effect of ROI was examined by collapsing activation across switch type for each ROI (Figure 3c). Activation increased along an anterior axis for some ROIs, with pre-PMd activation significantly greater than in PMd (paired t-test: t_14_=3.286, p=0.011, CI=0.27 to 1.28, *d*=0.85) and IFS activation showed a larger increase, in terms of effect size, relative to PMd (t_14_=3.733, p=0.007, CI=0.44 to 1.61, *d*=0.96). Activation in pre-PMd and IFS did not differ (t_14_=−1.175, p=0.312, CI=−0.71 to 0.21, *d*=0.30). However, activation in IFS was significantly greater than in aINS (t_14_=2.815, p=0.021, CI=0.16 to 1.16, *d*=0.73) and the aPFC (t_14_=6.494, p<0.001, CI=0.76 to 1.52, *d*=1.68) and activation in aPFC did not differ to the PMd (t_14_=0.359, p=0.725, CI=−0.58 to 0.82, *d*=0.09) (all p-values FDR-corrected). This provides some evidence of a posterior-anterior hierarchy for switching. However, this was for switching *generally*, and furthermore, the most anterior aPFC ROI was similar in activation level to the most posterior, PMd, ROI.

Activation was then collapsed across ROI to explore the main effect of switch type (Figure 3d). Comparison and side switch activation did not differ (paired t-test: t_14_=−0.393, p=0.700, CI=−0.55 to 0.38, d=0.10). Relative to side switching, activation for rule switching (t_14_=3.598, p=0.018, CI=0.35 to 1.38, d=0.93) and target switching (t_14_=3.040, p=0.027, CI=0.13 to 0.77, d=0.79) was significantly greater, and rule switching and target switching activation did not differ (t_14_=1.840, p=0.173, CI=−0.07 to 0.88, d=0.48). Relative to target switching, dimension (t_14_=−0.944, p=0.542, CI=−0.43 to 0.17, d=0.24) or task switching (t_14_=−0.532, p=0.700, CI=−0.51 to 0.30, d=0.14) activation did not differ (all p-values FDR-corrected).

### Supplemental voxelwise analysis of cognitive demands does not reveal an abstraction hierarchy or selectivity in PFC

We ran an unconstrained voxelwise analysis that directly tested the effect of cognitive demand, indexed by RT, on activation (Figure 3e). This did not render activation increases in frontopolar cortices for more demanding switches (e.g., dimension & task switching). Nor did task-switching, the most difficult switch, selectively activate any region within the frontopolar cortex, when contrasted with the easiest condition, comparison switching. Instead, areas of suprathreshold activation increase were distributed within a left lateralized set of regions comprising areas of multiple-demand cortex and the dorsal visual streams. The inverse contrast, did not render any regions of greater activation for less difficult switches at the cluster-corrected threshold.

### Cascade and rule abstraction ROIs show no evidence of an abstraction hierarchy in PFC

As a final step, we used the exact landmarks outlined by the cascade and rule theories to test for evidence of a functional hierarchy. 5mm spheres, centred on the ROI coordinates, were used to extract activation beta estimates from each switching condition and subject, and modelled in a ROI * switch type rm-anova. The cascade model ROIs (inferior frontal sulcus (IFS), dorsal premotor (PMd), and prePMd) showed a main effect of switching condition (F(5, 70)=6.275, p=0.001, 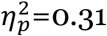), and no main effect of ROI (F(2, 28)=0.622, p=0.527, 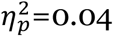) or significant interaction (F(10, 140)=1.500, p=0.201, 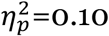) (p-values Greenhouse-Geisser adjusted). The rule abstraction ROI model (aPFC, IFS, PMd, prePMd) showed a main effect of switching condition (F(5, 70)=4.481, p=0.009, 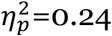), ROI (3, 42)=13.095, p<0.001, 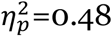) and no significant interaction (F(15, 210)=1.754, 0.122, 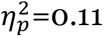) (all p-values Greenhouse-Geisser adjusted).

## Discussion

The aim of this study was to test predictions from alternative perspectives on human frontal lobe functional organisation. Neither perspective fully predicted the observed results.

A novel feature of the task design was that switches at different levels of abstraction were partially differentiated from general difficulty, as gauged by the response time costs. From the perspective of models that assert that the functions of the frontal lobes are organised hierarchically along a posterior-anterior abstraction axis (Koechlin et al. 2003; Badre and D’Esposito 2007), we predicted that switching costs associated with changes to rule mappings should activate more anterior frontal regions than those associated with stimulus dimension or target switches. We also predicted that task switches should specifically activate the most anterior PFC region (Koechlin et al. 1999; Dove et al. 2000; Rushworth et al. 2002; Braver et al. 2003). We could find no evidence to support a posterior-anterior functional axis. We also found no evidence that the aPFC plays a unique or specialised role in attentional switching, despite this being a prevailing view in the literature.

Instead, our findings demonstrate that cognitive control, responsible for the reconfiguration of attentional resources during switching, engages highly distributed neural systems. More specifically, switching-associated activation was evident throughout frontoparietal, subcortical and cerebellar regions. This included aPFC, even at the lowest levels of switching demand. This highly distributed pattern replicates several task-switching studies (Hampshire and Owen 2006; Cusack et al. 2010; Kim et al. 2012; Daws et al. 2020) and accords poorly with the notion that specific cognitive control processes are localisable to specific regions, including the tight mapping of switching to frontopolar cortex, as has been previously proposed (Koechlin et al., 1999; Rushworth et al., 2002; Braver et al., 2003).

Conversely, the multiple-demand perspective in its original formulation does predict that there should be patterns of distributed activation during switching conditions, in general (Duncan and Owen 2000). However, it also has been proposed that the extent of this system is flexible and will recruit additional neural resources, typically anterior frontal regions, in response to general increases in cognitive demand (Duncan, 2013; Crittenden and Duncan, 2014).

This failure to validate either prediction presents an intriguing conundrum. A wealth of imaging studies have reported functional dissociations within the brain volume that comprises multiple demand cortex (Christoff et al. 2001; Ullsperger and von Cramon 2001; Hampshire et al. 2007; Parkin et al. 2015; Crittenden et al. 2016; Mineroff et al. 2018). In many cases those dissociations have been robustly reproduced. The latter refinement of the MD perspective provides an explanation of why such dissociations might be evident. Therefore, it differs from hierarchical models as the increased anterior spread of task related activity within the PFC is predicted to be general for cognitive demands, irrespective of their abstraction (Jiang and Kanwisher 2003; Crittenden and Duncan 2014). The fact that no such anterior spread at higher switching demand was evident in the current study accords poorly with that explanation.

Indeed, for either of these accounts to be accurate, they must describe fundamental properties that generalise across contexts. When taken in combination, the results presented here and in previous studies accord with the notion that functional dissociations do exist within the lateral PFC, and that there may even be some level of hierarchy in network dynamics or regional function. However, the notion of an axis with concrete-abstract processing may be overspecified as it is based on results that do not generalise to tasks that probe related processes using somewhat different designs.

Arguably, this study did provide some evidence of a functional dissociation across lateral PFC regions. For example, the watershed transform parcelated the task-evoked PFC activation into regions centered on distinct peak foci, and those ROIs aligned well with areas previously reported to functionally dissociate in hierarchical models (Koechlin et al. 2003; Badre and D’Esposito 2007). Furthermore, those regions showed differential sensitivity to switching. However, the difference in activation presented as an inverted U-shape along the posterior-anterior axis, with activation in the most anterior ROI (aPFC) similar to the most posterior ROI (PMd). Moreover, activation uniformly increased in all PFC regions with cognitive demand and regardless of abstraction.

A more accurate account of the data from this study comes from the multiple-demand perspective as originally couched, or relatedly, the global workspace model (Dehaene, Kerszberg and Changeux, 1998). However, when taking a broader perspective, the challenge is to determine the functional basis of previously observed dissociations, and possible hierarchy, within MDC.

We propose several possibilities for future investigation. One possible explanation is that the apparent spread of activity with increased difficulty is a variant on the “imager’s fallacy” (de Hollander *et al.*, 2014). Specifically, application of a binary threshold to statistical maps can produce the impression of functional dissociations. Here, anterior PFC voxels were somewhat less active for switching in general, but activity increased with general difficulty. Although the latter effect was uniform across ROIs, if more anterior regions were just below and more posterior regions just above the statistical threshold at the lower switching level, this could lead to the illusion of a selective sensitivity of anterior region under higher switching demand. This same issue holds true for the notion of an increased spatial spread of MD with difficulty. This makes particular sense if MD is itself not considered to have a hard boundary, but instead, to capture a gradient of domain generality, where each brain region or voxel has some varying level of involvement across task contexts.

This latter notion of domain generality accords well with the observation of functional dissociations within MD, given the capacity of multivariate machine learning algorithms to identify tasks based on both activation and connectivity patterns therein (Crittenden, Mitchell and Duncan, 2016; Soreq, Leech and Hampshire, 2019). Relatedly, we have argued that it may be inappropriate to impose strong functional dissociations on what may essentially be a multivariate system that supports tasks through a many-to-many mapping system (Hampshire and Sharp, 2015; Lorenz, Hampshire and Leech, 2017). Such a relationship would explain discrepancies in mappings from studies using similar but non-identical task designs, that is due to the imposition of overly narrow experimental designs and hypotheses based on narrow sampling of task designs (Lorenz, Hampshire and Leech, 2017).

A further potential explanation relates to a series of studies where we have reported reproducible evidence of functional hierarchies across the frontal cortices, in terms of connectivity and regional activation analyses (Erika-Florence, Leech and Hampshire, 2014; Parkin *et al.*, 2015; Hampshire *et al.*, 2016, 2019). These studies highlight that brain regions show changes in activation patterns and dynamics even when performing the same task but that are dependent on different stages of learning. Specifically, as individuals practise novel tasks their response speed increases and trial-evoked activation within PFC steadily decreases along an anterior-posterior axis. Taken together, these “learning curves” demonstrate aPFC recruitment during demanding contexts that can be transient as individual’s behaviours become optimised and, consequently, global cognitive demand falls.

From this, we speculate that within-task learning occurs at different rates for more and less complex task manipulations, and if not accounted for, and an average of all trials of a type is taken, that could produce the illusion of greater aPFC activity under more complex conditions, when differential placement on the learning curve may underlie such differences.

It is important to address the present methodological approach and determine if this precluded us from observing evidence of a hierarchy or multiple-demand dissociation. First, it could be argued that our ML switching paradigm provided switching-magnitudes and variability across switch types that were insufficient to detect switching-specific aPFC activation or differences between PFC regions that would suggest a hierarchy. We consider false-negatives to be unlikely in this context as behavioural performance showed a significant variability across switch types. The comparison of the switches with low (side switch) vs high (task-switching) abstraction showed a substantial ~700ms difference in RT. Furthermore, there was sufficient power to detect significant differences in aPFC activation for multiple contrasts; however, these same contrasts were also evident elsewhere.

Second, our comparison of activation across the PFC was conducted using ROIs that were constructed in a data-driven manner. Importantly, these ROIs showed a high correspondence with, and contained, coordinates previously defined by the cascade (Koechlin, Ody and Kouneiher, 2003) and abstraction hierarchy theories (Badre and D’Esposito, 2007). However, to abate potential concerns that the focused ROI analysis was biased, we performed an unconstrained whole-brain analysis which tested the effect of switching-abstraction on voxelwise activation. Both contrasts supplemented and extended the ROI analysis by demonstrating that switching-abstraction, and direct comparisons between the least and most challenging switches (task>comparison), increased activation across PFC and beyond in bilateral occipital cortices dorsal visual streams. Strikingly, this absence of evidence in support of a functional hierarchy was also the case when directly measuring activation from the exact MNI coordinates previously defined by both hierarchical models.

Finally, it could be argued that activation increases for more abstract switches are explainable by these switches being more challenging and therefore requiring individuals to attend to stimuli for longer periods. This is unlikely to be the case here, as we used individuals trial RT to model activation per unit time, which essentially factors out activation differences due to visual-attentional processing times. Therefore, it is the case that even taking into account those differences, uniform increases in activation *per unit time* were evident for more difficult switching conditions.

## Conclusions

In summary, our results challenge predictions of prominent theories of abstraction and multiple-demand regarding the functional organisation of the frontal lobes. We demonstrate that neither of these models fully accounts for the distributed and uniform activation increases in the frontal lobes during switches with increasing abstraction and cognitive demands. However, the MD perspective in its original form is closer to the mark when explaining the results reported here. Further work is required to better specify a fundamental organisation of the lateral PFC that can be demonstrated to generalise across task contexts.

## Data and Code Availability Statement

Individuals preprocessed fMRI and raw behavioural data will be made publicly available upon acceptance for publication (via openNeuro.org).

## CRediT authorship contribution statement

**Richard Daws:** Formal analysis, Methodology, Visualization, Data curation, Writing - Original Draft. **Eyal Soreq:** Methodology, Visualization, Writing - Original Draft. **Yuqi Li:** Writing - Original Draft. **Stefano Sandrone:** Writing - Original Draft. **Adam Hampshire:** Conceptualization, Methodology, Writing - Original Draft, Investigation, Resources, Software, Supervision, Project administration, Funding acquisition.

## Acknowledgements

This work was supported by funding to Richard E. Daws from the Engineering and Physical Sciences Research Council (EPSRC) via the centre for doctoral training (CDT) Neurotechnology programme at Imperial College London. The funders had no role in study design, data collection and analysis, decision to publish or preparation of the manuscript.

## Notes

### Competing Interest Statement

The authors have declared no competing interest.

